# Type 1 diabetes genome-wide association analysis with imputation identifies five new risk regions

**DOI:** 10.1101/120022

**Authors:** Nicholas J. Cooper, Chris Wallace, Oliver Burren, Antony Cutler, Neil Walker, John A. Todd

**Author notes:** Corresponding authors: Nicholas Cooper and John Todd.

## Abstract

Type 1 diabetes genotype datasets have undergone several well powered genome wide analysis studies (GWAS), identifying 57 associated regions at the time of analysis. There are still many regions of smaller effect size or low frequency left to discover, and better exploitation of existing type 1 diabetes cohorts with meta analysis and imputation can precede the acquisition of new or larger cohorts. An existing dataset of 5,913 case and 8,828 control samples was analysed using genome-wide microarrays (*Affymetrix GeneChip 500K* and *Illumina Infinium 550K*) with imputation via *IMPUTE2* with the 1000 Genomes Project (phase 3) reference panel. Genotyping coverage was doubled in known association regions, and increased by four fold in other regions compared to previous studies. Our analysis resulted in new index variants for 17/57 regions, an expanded set of plausible candidate SNPs for 17 regions, and five novel type 1 diabetes association regions at 1p31.3, 1q24.3, 1q31.2, 2q11.2 and 11q12.2. Candidate genes for the new loci included *ITGB3BP, FASLG, RGS1, AFF3* and *CD5/CD6*. Further prioritisation of causal genes and causal variants will require detailed RNA and protein expression studies, in conjunction with genome annotation studies including analysis of physical promoter-enhancer interactions.

## 1 Introduction

Type 1 diabetes genotype associations have been explored through several well powered genome wide analysis studies (GWAS), identifying 57 associated regions (catalogued by ImmunoBase [10]). There are still many regions of smaller effect size or low frequency left to discover, and better exploitation of existing type 1 diabetes cohorts with more powerful and targeted methods is an informative step forward.

To declare a disease association region a SNP association should be observed with a P value less than the genome wide significance threshold (GWST) of 5 *\** 10^−8^ [17, 27]. Additionally it has been argued that any variants in a GWAS with *p <* 10^−5^ and *p <* 5 *\** 10^−8^ in another cohort can be use to define *pleiotropic* association regions [20]. Six of the 57 known regions have been declared using pleiotropic criteria and the remainder showed at least one variant with *p <* 5 *\** 10^−8^.

Given the huge effort required to acquire new cohorts, attempts have been made to gain maximum statistical power from existing datasets. Genome wide *imputation* of unmeasured SNPs [35, 37] is now routinely conducted for large genome-wide datasets and for type 1 diabetes has been performed several times to obtain datasets of up to 2.6 million SNPs[9, 4, 51] using *HapMap2* [28]. Simulations have shown that imputation can increase the power of a GWAS by 10% [45].

Reference panels used for imputation serve as large libraries of haplotypes to which we match our sparsely genotyped samples. Larger panels provide longer and more accurate matched haplotypes [35]. An ethnically diverse panel is advantageous, even for imputing for a European sample, because almost everyone has a small number of exotic haplotype segments [25].

The latest 1000 Genomes Project phase 3 haplotypes dataset was released during October 2014. This updated reference contains 79 million SNPs and 3 million other variants for 2,504 samples from 26 ethnic populations. The majority are monomorphic in Europeans, the remainder comprise ~15 million rare variants and ~10 million SNPs with MAF > 1% (which is a threshold below which imputation is no longer robust). Table 1 provides a breakdown of comparative densities between this reference set, *ImmunoChip*, and the GWAS microarray platforms used in this study.

**Table 1.**
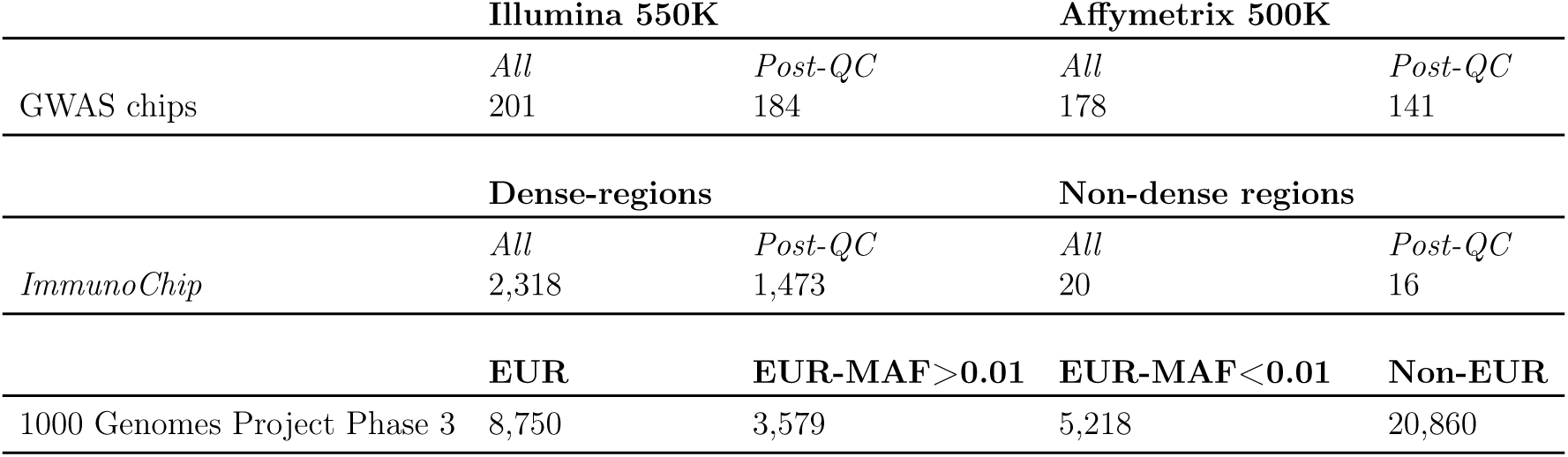
Table showing densities (SNPs per megabase) for GWAS chips versus *ImmunoChip*, versus the 1000 Genomes Project. Densities were calculated as medians to prevent skewness due to genome annotation gaps. The densities for the two GWAS chips and *ImmunoChip* were calculated with the full complement of SNPs and the post-QC set. Note that there is only roughly 20% overlap between SNPs in the two GWAS chips so the coverage for the Barret et al. study (8) would be near to the sum of the GWAS chip densities. *ImmunoChip* densities were calculated within dense autoimmune regions (which were specifically targeted to be as complete as possible as part of the design of the chip) versus the remainder of the genome. EUR = SNPs with MAF greater than zero in Europeans.

New associated regions have required increasing sample sizes to detect (Supplementary Figure 1), with power often supplemented by targeted genotyping of SNPs that fell marginally short of the GWST in the first pass. With the recent type 1 diabetes *ImmunoChip* study [40], new associations were facilitated through increased coverage in high-prior genomic regions. Based on Table 1, imputation with 1000 Genomes Project Phase 3 will more than double the *ImmunoChip* coverage in densely mapped regions, and should provide a four fold improvement in coverage for non-dense regions versus previous imputation analyses of type 1 diabetes datasets [9, 51].

Building on the success of *ImmunoChip* for investigating novel type 1 diabetes regions [40], candidate causal genes and SNPs, a similar analysis strategy has been implemented, now with an imputed genome-wide dataset.

*ImmunoChip* was designed to target regions of prior autoimmune association in greater detail, and due to its lower cost facilitated collection of large samples allowing comprehensive mapping of candidate causal variants in established regions. However, type 1 diabetes associations are also likely to exist outside of established autoimmune regions. By densely imputing from genome-wide arrays the opportunity arises to find SNPs in regions where LD between GWAS SNPs and the causal variants was too weak to create a GWST association.

## 2 Methods

### 2.1 Platform

The original set of genotypes used in this analysis were derived from the Affymetrix GeneChip Mapping 500K Set and Illumina 550K Infinium microarray platforms. Further detail for the data acquisition is provided in the original study [4].

### 2.2 Subjects

Samples, totalling 5,913 cases and 8,828 controls passing QC, for this study were derived from several sources. Supplementary Table 5 provides a summary of the sample characteristics and sources. The data sources have a complex structure with intersecting sets of markers common to both platforms and intersecting samples acquired using both platforms, this structure is described in Supplementary Figure 2.

The Affymetrix dataset originated from the WTCCC GWAS [11] and UKBS. The WTCCC data included controls and Bipolar disorder samples, the latter of which were used as extra controls as they showed very similar allele frequencies to controls [11]. The original study where this dataset was first used [4] also contained 3,305 USA Affymetrix samples from GoKinD/NIMH [13] which had lower data quality and and a subtly different structure of ethnicity.

The Illumina dataset was genotyped for use in the Barrett et al [4] study taking samples from the 1958 Birth Cohort (1958BC, N=6,929, [42]) as well as type 1 diabetes and control samples from the UK Genetic Resource Investigating Diabetes (UK GRID) cohort [49]. There were 1,444 1958BC samples typed on both platforms. For the 75,000 SNPs common to both platforms genotype data from the Affymetrix platform was used, and for imputation these samples were imputed alongside the uniquely Affymetrix samples. This decision was made in the interest of more closely balancing the size of the Affymetrix versus Illumina data sources, and to more closely following the cohort groupings in the original Barrett et al [4] study.

### 2.3 Family data

Data from Onengut-Gumuscu et al [40] collected using *ImmunoChip* [14] was available to us, but the cases and controls were largely overlapping with the current dataset. The family samples were independent, and were used as a reliability check for new associations and for meta-analysis with three imputed SNPs shared with the *ImmunoChip*.

### 2.4 Pre-imputation QC steps

#### 2.4.1 Genotype QC

Quality control of the original dataset was performed for a previous study [51]. Samples were excluded for non-European ancestry, both self report or PCA-inferred, or if they were duplicated or strongly related to another sample in the study. SNPs were excluded for MAF below 1% in cases or controls, or for HWE (*P <* 5.7 *\** 10^−7^), or for call rate below 95%.

#### 2.4.2 Alignment of datasets

Datasets were aligned to one another and then to the European groups of the 1000 Genomes Project phase III reference using the *annotSnpStats* R package (https://github.com/chr1swallace/annotSnpStats). Alignment used allele codes where possible. Allele codes are not helpful for ambiguous A/T or C/G genotypes, so in these cases alleles were aligned using reference allele frequency. Any SNPs with ambiguous allele codes and frequencies within 45% - 55% were considered too difficult to align and excluded. Alignment success of study datasets is shown in Supplementary Figure 10.

#### 2.4.3 Principal components and ancestry

To confirm that population structure had been adequately addressed, PCA was conducted for ancestry, both projecting from the 1000 Genomes Project samples (Supplementary Figure 3) and within the study dataset (Supplementary Figure 4). Details of the strategy used for dealing with ancestry effects are described in Supplementary Methods 1.4. Our resulting overall inflation *λ*_1000_ statistic was 1.02, where this is defined as the GC inflation lambda scaled for 1000 cases and 1000 controls [22].

### 2.5 Replication analysis of the Barrett et al GWAS

As an initial validation step the Barrett et al [4] GWAS was replicated with results shown in Supplementary Table 6. The correlation between the original and replicated Log10 P values was 0.988, where a non-perfect relationship was expected due to the absence of the USA samples in our reanalysis.

### 2.6 Imputation using *IMPUTE2*

Imputation of unmeasured genotypes was performed using *IMPUTE2* [37] with the recently released 1000 Genomes Project phase III cohort as a reference.

Pre-phasing [24] was not utilised, which slowed processing but should maximise accuracy. It has been suggested that controls imputed on one platform should not be compared to cases imputed on another [3]. Testing on chromosome 20 showed that imputing separately by genotyping chip reduced the *λ*_1000_ inflation estimate to 1.02, from 1.14 when imputed together. Further details of imputation methods are provided in Supplementary Methods 1.2.

A filter removing SNPs with MAF <1% was used and greatly speeds imputation as only 12% of the total reference set of SNPs are attempted. Furthermore, rare imputed SNPs are very prone to artifact and based on a prior imputation project [38], we found that the vast majority of extreme P-value false positives were for low MAF variants (< 1%). Additionally, given that recent GWAS studies that have discovered new loci for type 1 diabetes have had larger samples than the present analysis, we are already working at a power disadvantage, so our power to detect association at rare SNPs would be very low.

### 2.7 Analysis of uncertain genotypes

Imputation algorithms output uncertain genotypes, so rather than a sample being assigned a fixed genotype, they will be assigned a vector of probabilities. These uncertainties were analysed using *SNPTEST* using expectation maximisation (EM) to estimate parameters in the missing data likelihood [37]. To combine beta coefficients (log odds ratios) and P values from separate analyses of the Affymetrix and Illumina cohorts, meta analysis was used with weights proportional to the inverse of the variance (see Supplementary Methods 1.5) for more details. For Bayesian analyses we simply summed the Log10 Bayes Factors for the two sources.

One of the most useful QC statistics for evaluating imputation success are the Information (‘info’) scores (for further details see Supplementary Methods 1.3). Information scores were higher for the Illumina data with a median score of 0.95 versus 0.89 for Affymetrix. Various studies have used different cutoffs for info scores, including 0.3, 0.5, and 0.8 [3]. Here we used 0.25 as failure threshold and less than 0.75 as a warning threshold.

#### 2.7.1 Imputation quality control

In order to minimise the effect of artifactual imputation, various quality control (QC) checks were performed for each SNP. Some checks were considered as potential ‘warnings’ while others were considered pass or fail. See Supplementary Methods 1.10 for the specific criteria.

Using these criteria, 10% of imputed SNPs failed, 42% passed with no warnings, 37% had one warning, 9% had two warnings, the remaining 3% had three or more warnings. The warnings are now stored with the summary statistics for the dataset so that future analyses can make the QC stringent or relaxed according to their application. The failing SNPs from all analyses were excluded, while ‘warnings’ were used to inform interpretation of any significant SNP association results.

#### 2.7.2 Index SNPs, conditional analysis and colocalisation

The index SNP, conditional signal, and colocalisation analyses were conducted as per the recent ImmunoChip study [40]. These methods are also described explicitly in Supplementary Methods 1.6-1.7. Conditional analyses were also run with *SNPTEST,* and a scan for pleiotropic associations showing *P <* 10^−5^ where *p <* 5 *\** 10^−8^ has been observed in another autoimmune disease, was undertaken.

#### 2.7.3 Credible Sets

Bayes factors were generated directly by using *SNPTEST* with ‘bayesian’ analysis using the ‘score’ method. Log10 Bayes factors from the Affymetrix and Illumina datasets were combined using a simple sum to obtain meta analysis Bayes factors [23]. Further details are provided in Supplementary Methods 1.8. The cumulative sum of the sorted posterior probabilities for all SNPs in each region were used to define 99% credible sets.

## 3 Results

### 3.1 Immune and type 1 diabetes association regions replication

### 3.2 New type 1 diabetes regions

Six new associations in five novel regions are presented in Table 2. These five regions are visualised in Supplementary Figures 12-14. Converging patterns of LD based association in these plots, including for directly genotyped SNPs from our Barrett et al [4] replication and from the *ImmunoChip* study [40], show that these associations are not due to extreme random or isolated imputation artifacts.

**Table 2.** New type 1 diabetes regions identified by this analysis. The regions either satisfy the GWST or satisfy the pleiotropic threshold of *p <* 10^−5^, when another related autoimmune condition has a GWST result for the SNP. Regions are defined as +/- 0.1 cM from the Index SNP. Index SNP locations are available in Table 4, except for rs12068671 which is at chr1:172,681,031. Alleles are major>minor and ORs are >1 when cases have a higher frequency of the minor allele. The variant, rs70940197, is an indel. ‘Info’ scores (range 0-1) are standard output from *IMPUTE2* and *SNPTEST* and reflect the amount of information available for imputation, increasing with the number of SNPs available highly correlated with the SNP being estimated. MAF is the minor allele frequency in Europeans in the 1000 Genomes Project legend file. P values for coeliac were used to satisfy a posterior probability > 0.9 and came from an *ImmunoChip* study [50]. Co-localisation was examined for all *ImmunoBase* catalogued immune diseases, but both co-localised hits related to the same coeliac dataset. rs2269242* is in full LD with rs2269240 in this dataset (position=64,109,264, C>T). † rs118000057 was confirmed using meta analysis with the type 1 diabetes family dataset from Onengut-Gumuscu et al [40] (it was one of three top novel regional SNPs with potential for such). The imputation analysis P value was 7.11 *\** 10^−7^, whilst the family TDT P value was 0.007.

| Location | Range: Index +/- 1 cM |  |  | Index SNP | OR | P value | Alleles | Info | MAF | P (Coeliac) |
| --- | --- | --- | --- | --- | --- | --- | --- | --- | --- | --- |
| 1p31.3 | 64,102,590 | - | 64,168,032 | rs2269242* | 1.18 | $2.14 * 10^{-8}$ | G>A | 0.98 | 0.24 | |
| 1q24.3 | 172,648,956 | - | 172,871,189 | rs78037977 | 0.78 | $3.06 * 10^{-9}$ | A>G | 0.83 | 0.12 | |
| 1q24.3 | 172,648,956 | - | 172,871,189 | rs12068671 | 0.85 | $7.92 * 10^{-7}$ | T>C | 0.97 | 0.20 | $1.40 * 10^{-10}$ |
| 1q31.2 | 192,469,607 | - | 192,547,791 | rs10801121 | 1.14 | $5.74 * 10^{-6}$ | A>G | 0.90 | 0.29 | $8.82 * 10^{-12}$ |
| 2q11.2 | 100,544,267 | - | 100,969,443 | rs70940197 | 0.86 | $1.00 * 10^{-8}$ | G>GTGATGA | 0.97 | 0.37 | |
| 11q12.2 | 60,680,130 | - | 60,856,960 | rs118000057 | 1.51 | $\dagger 1.24 * 10^{-8}$ | T>G | 0.71 | 0.02 | |

Three novel regions, 1p31.3, 1q24.3 and 2q11.2, satisfied a GWST in the primary analysis. Within 1p31.3, two index SNPs had equal P values: rs2269242 and rs2269240. The latter was *P <* 10^−6^ in the *ImmunoChip* analysis, and the nearby variant rs2269241 was marginally short of satisfying the GWST in the original Barrett et al study [2009].

The index SNP rs78037977 for 1q24.3 was also proposed as a new type 1 diabetes locus via Bayesian colocalization analysis for shared controls in a recent study [20]. Additionally, there was a further pleiotropic association in the same region with rs12068671. The adjusted OR covarying for UK region was slightly weaker at 0.805 suggesting slight inflation due to ancestry for this SNP. It should be noted that the converging study results mentioned here involved considerable overlap between subject cohorts.

A post-hoc meta-analysis was conducted on two SNPs known to be measured on ImmunoChip that fell marginally short of satisfying the GWST, allowing the 11q12.2 variant, rs118000057, to be included in Table 2.

#### 3.2.1 Regions satisfying a pleiotropic threshold

The current list of 867 SNPs with *P <* 5 *\** 10^−8^ in any *ImmunoBase* disease was downloaded and cross referenced to the results of this study. From this set, 91 SNPs were at or below GWST in this study, and 144 were at *P <* 10^−5^. Two SNPs from novel regions satisfied this condition, rs12068671 from the same association region as rs78037977 above, and rs10801121 from 1q31.2. Both SNPs were associated with coeliac disease with *P <* 10^−9^ [50].

#### 3.2.2 Comparison to ***ImmunoChip results***

Four of six novel index SNPs, rs78037977, rs12068671, rs10801121, and rs118000057, were directly genotyped on *ImmunoChip* and tested previously [40]. The latter SNP was confirmed using meta-analysis with the *ImmunoChip* families data. For the first four SNPs listed, the P values were higher and the ORs were considerably weaker in the family dataset at 1.06, 0.94, 0.95 and 0.96 respectively. The OR and significance statistics were more similar in the case-control dataset, but this dataset is not independent, using most of the same type 1 diabetes subjects. None of these SNPs had very poor signal clouds in the *ImmunoChip* dataset, with only rs78037977 showing some small clustering bias. Only rs2269242 and rs70940197 were unique to the imputation dataset.

#### 3.2.3 Cross checking of results

To increase confidence in the validity of results some checks of assumptions were carried out: (i) associations were confirmed as null on the Bipolar disease cohort to ensure that their use as controls was justified; (ii) OR and P value statistics were compared against models that included region and sex covariates showing that ORs were not reduced; (iii) GWST regions were confirmed not to fall within a list of cytobands with a strong association for UK geographical regions; and (iv) failures and warnings were generated based on post imputation QC metrics that excluded inconsistent findings between Illumina and Affymetrix datasets.

### 3.3 Conditional analyses

Conditional analysis was run for regions with a result passing the GWST. Results for regions with conditional associations from the *ImmunoChip* study [40], and any additional results passing a region-wise or global conditional Bonferroni correction are presented in Table 3. Three from five regions with conditional signals in the *ImmunoChip* study were at the region-wise threshold, and the first additional signal for 10p15.1 satisfied the global threshold. There were two novel conditional signals, neither of which met the global corrected P value criteria.

**Table 3.** Results for conditional analyses. Conditional results satisfying a Bonferroni correction for the number of SNPs within each region, or attempted replications of conditional associations from the *ImmunoChip* study [40] are tabulated. Those satisfying an overall Bonferroni correction for ~37,000 conditional SNP tests *P <* 1.35 10^−6^ have a **bold** rs id. Replication attempts failing at both thresholds are presented below the middle line. ‘Reg.Size’ is the number of SNPs tested in each region, ‘info’ is the SNPTEST information score, ‘Allele’ shows the major>minor alleles. A X for ‘iChip’ means this region had a conditional hit in [40], ‘Repl’ is for when such a hit was replicated and ‘Novel’ is for regions with no previously reported conditional signals. The ‘Condition on’ column shows which index SNP, and/or first level conditional SNP, was conditioned on for each result.

| Location | SNP | Position | OR | P value | Reg.Size | info | Allele | MAF | iChip | Repl. | Novel | Condition on |
| --- | --- | --- | --- | --- | --- | --- | --- | --- | --- | --- | --- | --- |
| 2q24.2 | rs141102226 | 162,992,004 | 0.44 | $3.03 * 10^{-6}$ | 1,313 | 0.71 | A>T | 0.01 | ✓ | ✓ | | rs33998987 |
| 10p15.1 | **rs4749924** | 6,082,396 | 0.86 | $1.03 * 10^{-7}$ | 1,580 | 0.97 | A>C | 0.34 | ✓ | ✓ | | rs12722496 |
| 12q13.2 | rs184688715 | 56,358,737 | 1.61 | $3.49 * 10^{-5}$ | 1,260 | 0.64 | A>C | 0.03 | | | ✓ | rs7297175 |
| 19p13.2 | rs34536443 | 10,463,118 | 0.68 | $1.82 * 10^{-5}$ | 1,153 | 0.61 | G>C | 0.03 | ✓ | ✓ | | rs142155407 |
| 20p13 | rs202458 | 1,713,957 | 0.84 | $1.03 * 10^{-5}$ | 934 | 0.86 | G>T | 0.15 | | | ✓ | rs6043405 |
| 11p15.5 | rs7115640 | 2,194,914 | 1.13 | $3.96 * 10^{-4}$ | 1,103 | 0.62 | G>A | 0.35 | ✓ | | | rs689 |
| 18p11.21 | rs2542158 | 12,788,424 | 1.23 | $1.01 * 10^{-4}$ | 1,370 | 0.79 | G>A | 0.08 | ✓ | | | rs35153695 |
| 10p15.1 | rs41295159 | 6,148,535 | 0.53 | $5.53 * 10^{-5}$ | 1,579 | 0.67 | C>G | 0.01 | ✓ | | | rs12722496,<br>rs4749924 |

### 3.4 Updated credible sets for causal variants

Credible sets were generated for novel regions and those at a GWST reported previously. Table 4 shows separate and combined BFs and posterior probabilities for each index SNP. Three credible set counts were reduced while 17 were expanded, many of these considerably. Credible set sizes ranged from 1 to 247, where the median size was 2.8% of the number of SNPs in each region (including an extension of +/- 50 kb).

**Table 4.**
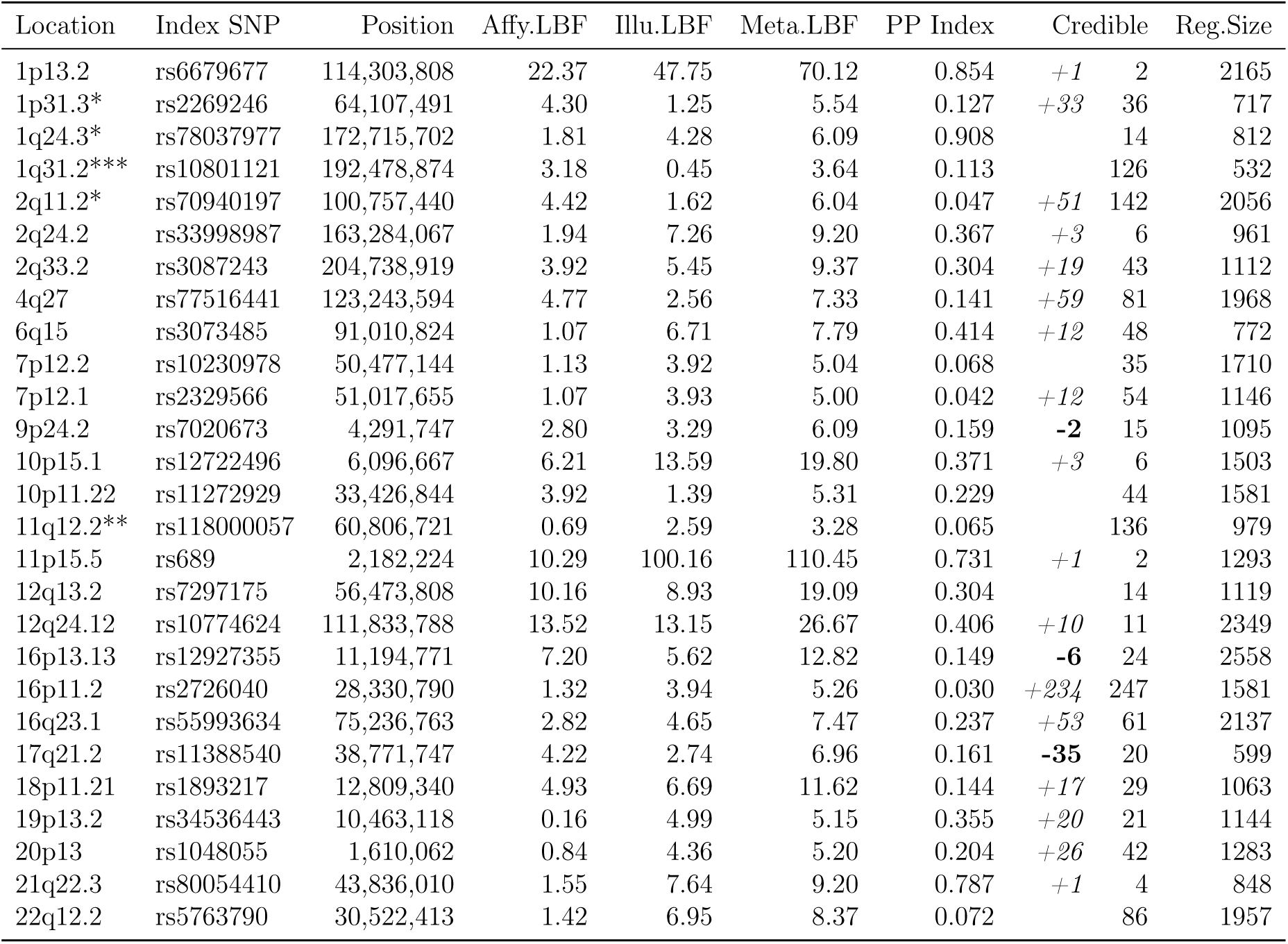
Credible SNP sets for associated regions. Credible sets for all type 1 diabetes regions passing GWST, novel type 1 diabetes regions passing GWST*, and novel type 1 diabetes regions confirmed using co-localisation*** or meta-analysis with the families dataset from Onengut-Gumuscu et al [40]**. ‘Affy’ and ‘Illu’ LBFs are the Log10 Bayes Factor (LBF) scores for each Index SNP, originating from SNPTEST Bayesian score tests. The meta-analysis LBF is simply the sum of the LBFs from the two subsets. ‘PP Index’ is the probability of the Index SNP being the causal variant, considering all other SNPs within the region as the alternative candidates. ‘Credible’ is the size of the 99% credible set. The parallel *+* and **-** numbers show the increase or decrease in the size of these sets versus the credible sets from the *ImmunoChip* paper. ‘Reg.Size is the number of SNPs in the association region (for novel hits +/1 1cM was used) +/-50 kb passing QC. The numbers in brackets next to the credible counts are the increase in the size of the set versus Onengut-Gumuscu et al [40].

Based on average densities, this imputed dataset should have at least twice as many SNPs in each region as *ImmunoChip*, and this was indeed the case with a mean set size of 48.2 for imputation versus 22.8 for *ImmunoChip*.

## 4 Discussion

Imputation has permitted the identification of five new type 1 diabetes risk regions at GWST and further fine mapping of established loci, bringing the total number to 62. New index SNPs were supported for 16 of these regions, and 17 credible sets were expanded versus those derived in the *ImmunoChip* paper [40].

To provide support for these findings, the validity of the imputed genotypes was confirmed in several ways. Assumptions regarding the suitability of the Bipolar disease cohort as controls were satisfied. Consideration of potential ancestry confounds and sensitivity testing with covariation for UK region supported the effect sizes detected in the primary models. Consistency of allele frequencies between Affymetrix and Illumina analyses, and for controls against European samples from the 1000 Genomes Project were also confirmed for reported associations.

Four of the five new regions have previously been associated with at least one other autoimmune or autoinflammatory disease.

The 1q24.3 region with index SNP rs78037977 was identified in a Bayesian colocalization analysis with shared controls [20]. A second SNP, rs12068671, was also confirmed using pleiotropic association with coeliac disease [50]. This region has also been associated for Crohn’s disease and ulcerative colitis (UC) [21, 31]. A previously proposed causal gene for this region is *FASLG* (Fas ligand TNF superfamily, member which has through interaction with its ligand FAS has a critical role in inducing apoptosis and cytotoxicity [46], although nearby tumour necrosis factor (TNFSF) genes, *TNFSF4* and *TNFSF18*, are also plausible candidates.

The 1p31.3 region was previously only just shy of the GWST [4]. The dual index SNPs rs2269242/rs2269240 fall within an intron of *PGM1* (phosphoglucomutase 1, glucose metabolism). This gene is not considered a good causal candidate for autoimmune diabetes. Another nearby gene with a potential functional connection is integrin beta 3 binding protein beta3-endonexin (*ITGB3BP*) which is involved in cell death signalling [43].

The 1q31.2 region was associated previously with type 1 diabetes utilising pleiotropy with coeliac disease for rs10801121 [50]. The region fell just short of the GWST in three previous type 1 diabetes studies [4, 9, 41], and also has a pleiotropic association with multiple sclerosis (MS) [29]. The proposed candidate gene in these previous studies is *RGS1* (regulator of G-protein signalling 1), which is involved in B cell activation and proliferation. Experimental work has shown RGS1 is expressed in T-cells in the intestinal intra-epithelial lymphocyte compartment [26], providing another potentially gut/microbiome linked type 1 diabetes region [44, 34, 1]. Until further analyses are performed linking the most associated variants with expression of certain genes in a region we need to be cautious assigning too much weight to a particular gene as being causal for disease. For example, chromosome conformation capture results in primary human blood cells indicate that TROVE2, which has been associated with the autoimmune disease, systemic lupus erythematosus, should also be considered a candidate causal gene in this region [30]. Data can be browsed at www.chicp.org, and the strongest weightings for this SNP were with macrophage and neutrophil tissue types.

The novel 2q11.2 locus was indexed by the insertion/deletion (indel) variant rs70940197 lying within an intron of the gene *AFF3* (AF4/FMR2 family member 3). This gene was identified as associating with rheumatoid arthritis (RA) [5] and Chinese systemic lupus erythematosus patients [12]. The G allele at rs10865035 was also linked to better anti-TNF treatment response in RA [47]. Weak evidence for this assoication with type 1 diabetes was found previously [49, 9] and both these studies posited *AFF3* as the causal gene. If *AFF3* is shown to be the causal gene, considering the likely causal candidates for 1q24.3 are also related to TNF, these results provide support for the involvement of this pathway, which has been linked to type 1 diabetes pathogenesis [8, 33, 6].

The 11q12.2 region was one of three non-type 1 diabetes *ImmunoChip* dense regions to achieve *P <* 10^−5^, so was followed up with additional meta-analysis using the type 1 diabetes families dataset [40], where the revised P value passed the GWST. The region has also been confirmed in MS, UC and Crohn’s disease [29, 31]. The index SNP was rs118000057, with nearby genes: prostaglandin D2 receptor 2 (*PTGDR2*), *CD5* and *CD6*. The most likely candidates are *CD5* and *CD6* which are expressed on the surface of T cells and are also implicated by chromosome conformation capture data [30] where the strongest weightings are linked to naive and activated CD4 cell types. Experiments have shown that the absence of CD5 increases the responsiveness of thymocytes to T cell antigen receptor in vitro [48], and that variants within CD6 may mediate response to TNF-alpha inhibitors in rheumatoid arthritis [32].

Fine mapping was facilitated in this study firstly through conditional analyses that identified regions likely to contain more than one causal SNP, and secondly through the creation of credible sets likely to contain the true variant for each region. Three from five conditional regions identified in the *ImmunoChip* study [40] were replicated at a region-wise P value, and the remaining two had P<0.001. Previously unreported conditional signals were detected at a region-wise P value for 12q13.2 (*IKZF4* [18]) and 20p13.

The total size of credible sets for regions passing the GWST was doubled versus Onengut-Gumuscu et al [40], reflecting the map density ratio of the 1000 Genomes Project versus *ImmunoChip*. Although most set counts increased, some of these may be due to reduced power attenuating the relative differences between top and middling candidates in some regions. A few credible SNP sets were reduced in size, potentially due to inclusion of some highly associated SNPs that were previously unmeasured.

Index SNPs reflect the best available candidate causal variant in each region. In contrast to the novel regions, 14 from 25 top SNPs from replicated type 1 diabetes regions were *not* measured in the *ImmunoChip* study *and* obtained a larger Bayes Factor than the corresponding *ImmunoChip* index SNP.

Our sample size was smaller than recent GWAS in type 1 diabetes and other autoimmune conditions (Supplementary Figure 15). We hoped greater coverage would provide new associations emerging from previously untyped variants. Four out of five novel regions had an index SNP typed on *ImmunoChip*, and several of these SNPs were P>0.05 and showed a weaker OR in the independent families dataset. These SNPs may have shown stronger effect size in the imputation analysis due to chance, displaying winners curse against previous findings. These specific variants may have benefitted from comparison to a substantially different control group, may have been subject to genotyping bias on *ImmunoChip*, or may show inflated association due to artifact in imputation. An alternative explanation is that because a portion of the family dataset have been selected with the goal of obtaining multiple affected siblings, these families may have a greater than average HLA burden of risk, weakening associations with variants of low effect size.

Our newly identified disease-associated SNPs may also have been tested in the Bradfield et al [9] analysis. However, the authors provided an explanation for lower than expected power in that study: the cohort composition did not allow use of British controls with British cases and American controls with American cases. Additional covariates for ancestry were needed to control for this and as a result power was lost despite using a larger sample size and imputing to a greater coverage than any previous type 1 diabetes analysis.

The conditional and credible set analyses had sufficient power to identify some potential causal SNPs that were previously untested. However, no *third* signals from the *ImmunoChip* analysis were replicated, and some credible sets were very large. While comprehensive, large credible sets provide limited scope to be refined via annotation to a usefully small set of variants. The lack of replication for some conditional signals from the *ImmunoChip* study [40] could reflect either a power deficit, or that with expanded coverage new variants are true causal SNPs that were previously tagged indirectly by multiple SNPs in LD. To better address these issues these results would benefit greatly from combination via meta analysis with an additional imputed cohort.

We conclude that genotyping of more type 1 diabetes samples with genome-wide imputation will allow significant expansion of the set of type 1 diabetes regions. MS and RA have similar a genetic risk to type 1 diabetes but have identified far more association regions. This not only serves as an example of the benefits of additional power, but may also provide a valuable resource for colocalisation analysis. The RA dataset has been imputed to 10 million SNPs [39], and considering the greater sample size, would be likely to provide pleiotropic support for ~10% of 80 regions from this study that showed *P <* 10^−5^. If imputed using the same reference, other large cohorts in MS and the inflammatory bowel diseases could provide similar utility.

## Author contributions

NJC carried out statistical analyses and wrote manuscript. NW prepared data and reviewed/edited manuscript. CW assisted in the analyses and reviewed/edited manuscript. JAT directed research, provided biological expertise and reviewed/edited the manuscript.

## Acknowledgements

We gratefully acknowledge all individuals who provided biological samples or data for this study. This research uses resources provided by the Type 1 Diabetes Genetics Consortium, a collaborative clinical study sponsored by the National Institute of Diabetes and Digestive and Kidney Diseases (NIDDK), the National Institute of Allergy and Infectious Diseases (NIAID), the National Human Genome Research Institute (NHGRI), the National Institute of Child Health and Human Development (NICHD) and JDRF and supported by grant U01 DK062418 from the US National Institutes of Health. Further support was provided by grants from the JDRF (9-2011-253 and 5SRA-2015-130-A-N) and the Wellcome Trust (091157 and 107212) to the Diabetes and Inflammation Laboratory at Oxford and Cambridge Universities. This study makes use of data generated by the Wellcome Trust Case Control Consortium, funded by Wellcome Trust award 076113; a full list of the investigators who contributed to the generation of the data is available from http://www.wtccc.org.uk/. CW is funded by the Wellcome Trust (WT107881) and the MRC (MC_UP_1302/5). John Todd is the guarantor of this work and, as such, had full access to all the data in the study and takes responsibility for the integrity of the data and the accuracy of the data analysis.

## Availability of data and materials

The dataset supporting the conclusions of this article is available in the ImmunoBase repository, [https://www.immunobase.org].

